# A peek into Western Indian Ocean microbial richness: a pilot for a coral microbiome study

**DOI:** 10.1101/829564

**Authors:** Sammy Wambua, Hadrien Gourlé, Etienne P. de Villiers, Joyce Ngoi, Angus Macdonald, Erik Bongcam-Rudloff, Santie de Villiers

## Abstract

Microbial communities are essential components of natural ecosystems. Of the global oceans, the Indian Ocean remains the least studied in terms of its microbial diversity, despite it being a highly dynamic tropical water body. Metagenomics methods have significantly advanced studies in marine microbial ecology in recent years. Preliminary metabarcoding assessments are recommended to mitigate against the associated costs, prior to the metagenomics study, to give an impression of the diversity expected and determine the sequencing effort required.

We report here the first metabarcoding survey of bacterial diversity of the western Indian Ocean (WIO) using samples used for optimizing environmental DNA (eDNA) isolation as pilot experiment for a metagenomic study investigating the coral-reef microbiome of the region. Sampling of water and sediment samples was done near-shore sublittoral and within the coral reef.

About 3,000 microbial ribotypes were inferred, from which 41 phyla were uncovered. Sediments registered higher alpha diversity than seawater samples. The phylum Proteobacteria was dominant with its members constituting over 60% of the ribosomal sequence variants (RSVs). The other abundant bacteria were members of Bacteroidetes and Cyanobacteria phyla. We identified bacterial species with potential broad biotechnological applications, underscoring the WIO’s richness and the usefulness of eDNA metabarcoding approaches in bioprospecting as well as monitoring and/or surveying marine ecosystems.

## Introduction

Microbial communities play central roles in ecosystem functioning. In marine environments microorganisms are thought to be vital for essential ecosystem processes such as decomposition of organic matter and nutrient cycling (Das et al., 2006). Furthermore, shifts in population composition of coral-associated microorganisms often correlate with the onset of diseases and coral-bleaching (Krediet et al., 2013), suggesting a likely role of microbes in coral health and, therefore, the stability of reef ecosystems. Besides health, microbes are also known to be critical for coral nutrition (Rosenberg et al., 2007) homeostasis and protection against diseases (Godoy-Vitorino et al., 2017).

These observations underscore the importance to characterize microorganisms and their ecological niches as a strategy to clarify their roles and the environmental conditions they respond to (Glasl et al., 2019). This understanding is especially critical currently in the face of increasing climatic and human-related threats on marine ecosystems leading to significant loss in biodiversity and ecosystem function. Conservation and management strategies therefore need rapid biodiversity survey tools (Deiner et al., 2017). Comprehensive understanding of microorganisms can be used as a proxy for such surveys and for predicting responses of marine ecosystems because microbial communities respond and quickly adapt to disturbance (Ainsworth et al., 2010). Recent advances in sequencing technologies and bioinformatics analysis offer more efficient approaches to study microbial ecology which, in turn, improved our appreciation of the importance of microbial communities in shaping the dynamics of diverse environments.

Direct genetic analysis from an environmental sample is increasingly becoming popular through metagenomic approaches. Besides population structure, metagenomic studies use genomes recovered from the environment to uncover and generate hypotheses on functional diversity (Thomas et al., 2012), which may be complemented and confirmed through metatranscriptome studies. Metabarcoding is a less expensive approach employed primarily for the identification of multiple species in an environmental sample often based on the diversity of one gene (Taberlet et al., 2012), such as the 16S rRNA gene for prokaryotic studies. To ensure credible findings, these approaches require sufficient high-quality nucleic acid representative of all cells present in an environment for library production and sequencing (Thomas et al., 2012). This is often complicated as there are numerous factors from origin, state and transportation from sampling sites, to primers and library preparation biases that influence the quality and quantity of nucleic acids. One main challenge to obtaining good environmental DNA from marine ecosystem is the risk of degradation due to dilution and effects of salinity, tides, currents and transportation (Deiner et al., 2017). For this reason, preliminary metabarcoding assessments are recommended prior to metagenomic studies (National Research Council, 2007) to provide insight into the microbial diversity expected and guide on the depth of metagenomic sequencing required.

Although WIO is believed to host the second hotspot of coral reef biodiversity globally (Obura et al., 2017), it remains understudied with reference regard to microbiome analysis because the region lacks the necessary technologies and expertise. To our knowledge no molecular-based baseline has been documented on the microbial diversity of the WIO.

This preliminary report previews the first description of coastal WIO microorganisms identified by sequencing 16S rRNA gene from seawater and sediment samples. These samples were used for the optimization of microbial DNA isolation for a metagenomic study aimed at profiling the coral reef microbiome of the same region.

## Method

Sampling was done in August 2016 at low tide in Kuruwitu conservancy, a locally managed marine area on the Kenyan coast of the WIO with a tidal range of about 3.9 meters. We sampled two sites, about 50 meters apart, within the reef flat: close to the shoreline where no corals were found (which we termed, “near-shore sub-littoral”), and further from the shoreline where corals were thriving (“coral reef”). Microbial DNA was isolated from four litres of seawater and 2.5 g sediment samples within two hours post-sampling using the PowerWater^®^ kit for seawater and PowerSoil^®^ kit (MoBio Laboratories) for sediment samples. The variable regions V2-4-8 of 16S ribosomal RNA gene were amplified by PCR using the Ion 16S™ Metagenomics Kit primer set, and libraries were prepared from the resulting amplicons with the Ion Plus™ Fragment Library Kit and sequencing done on an Ion PGM™ platform (ThermoFisher Scientific). Due to logistical constraints we were only able to sequence the water samples in duplicate.

The amplicon sequence data were analysed with the *DADA2* (version 1.13.0) bioinformatics pipeline (Callahan et al., 2016) in the R programming environment (version 3.6.1)by R Core Team (2019). Raw reads were trimmed to 300 base pairs and filtered to exclude reads with fewer than 100 base pairs. After dereplicating and carrying out error learning model, 2,915 ribosomal sequence variants (RSV) were inferred from the resulting reads. Taxonomy was assigned by exact matching of each RSV to the SILVA database (release 132) formatted for *DADA2* (Callahan, 2018). *DECIPHER* version 2.13.0 (Wright, 2016) and *phangorn* version 2.5.5 (Schliep, 2011) packages were used to construct a phylogenetic tree. The resulting RVS and taxa tables, phylogenetic tree and sample metadata were merged using *phyloseq* version 1.28.0 (McMurdie & Holmes, 2013) and used in subsequent microbial community analyses.

## Results

Across all samples, a total of 569,106 high-quality reads were generated from of the six samples’ amplicon sequences, from which a total of 2,915 microbial RSVs were inferred. A rarefaction analysis for the identified RSVs verified that all samples had been sequenced to sufficient depth to capture representative microbial diversity (Fig 1).

**Fig 1:**
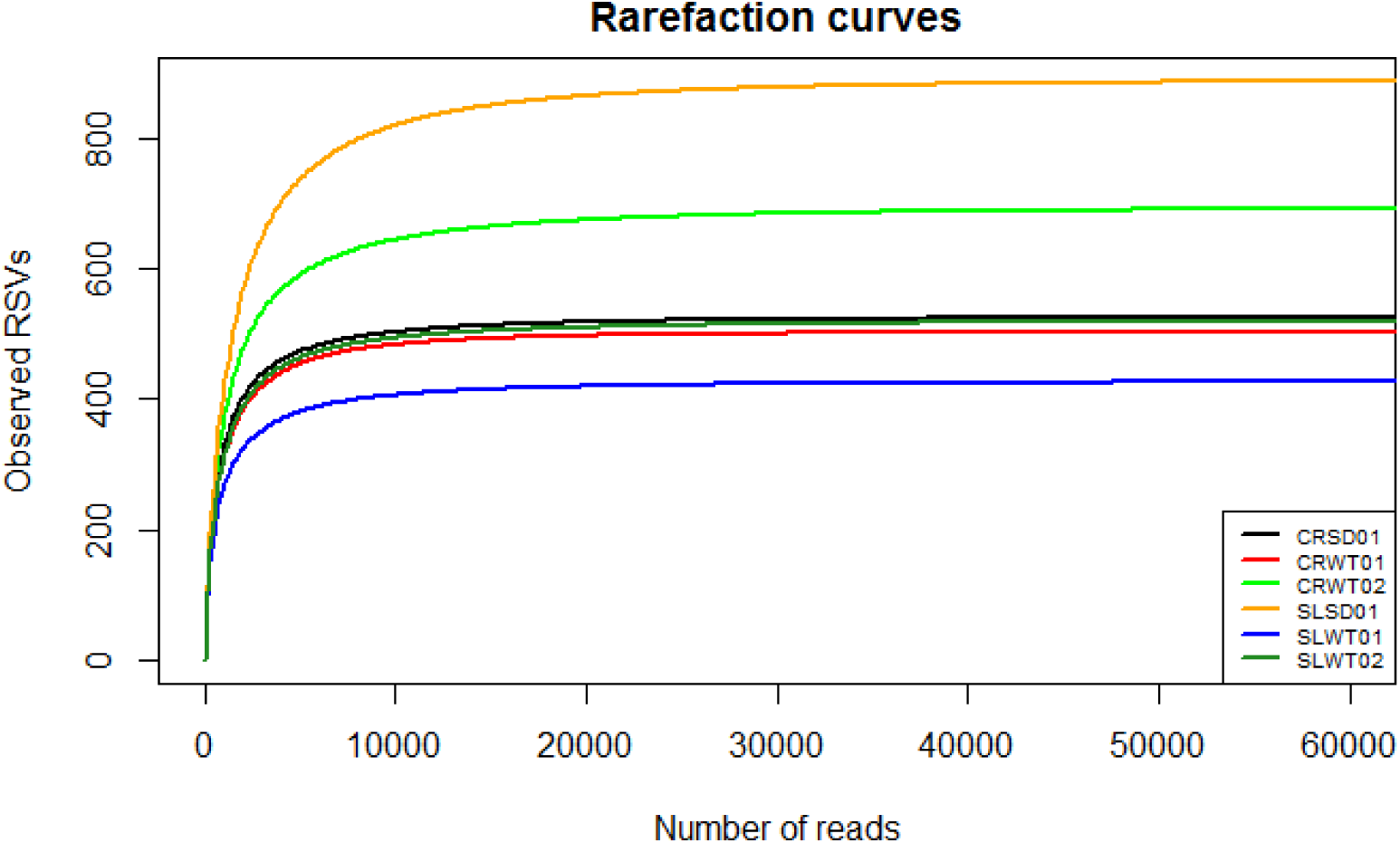
Rarefaction curves of bacterial RSV richness on every sample using the divisive amplicon denoising algorithm (DADA) approach. CR = coral reef location, SL= near-shore sublittoral location, SD = sediment, and WT = seawater.

Compared to seawater, sediment samples recorded higher alpha diversity metrics, calculated as observed richness (ANOVA *P =* 0.288) and Shannon diversity (ANOVA *P =* 0.228), which was not statistically significant (Fig 2). Community composition visualized through Principal coordinates analyses (PCoA) based on Bray-Curtis and Unweighted UniFrac distances revealed clustering of seawater samples. Sample type accounted for over 40% variance (Fig 3) although the differences between microbial population in seawater and sediment samples were not statistically significant as measured by permutational multi-variate ANOVA (PERMANOVA) on both Bray-Curtis (*P* = 0.067) and unweighted UniFrac (*P* = 0.067) dissimilarities. Of the top 30 most abundant genera, only five were represented in both water and sediment samples regardless of the sampling site (Fig 4).

**Fig 2:**
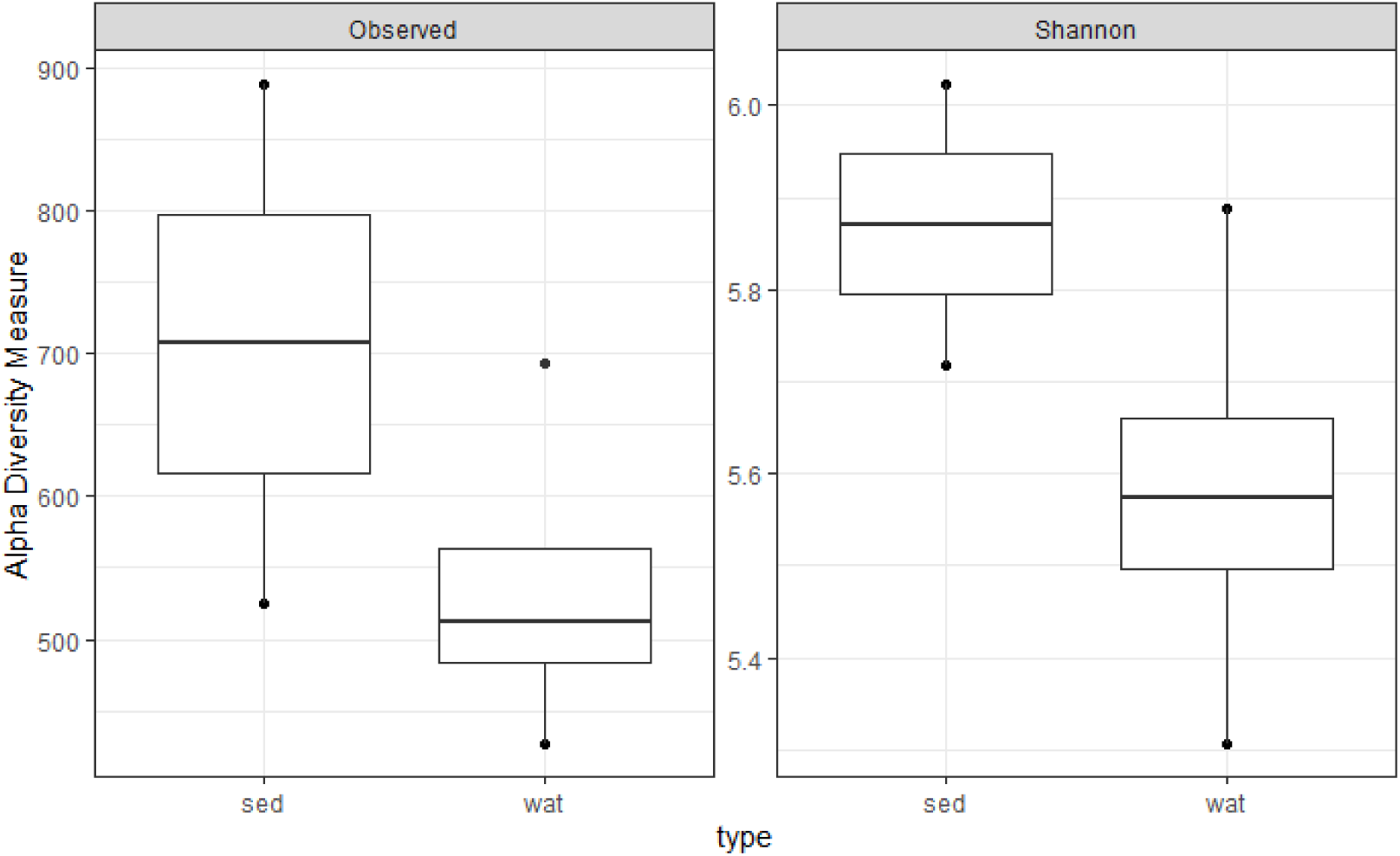
Plots of observed amplicon sequence variant (ASV) richness and Shannon diversity across samples, which are grouped by type: sed = sediments, wat = seawater.

**Fig. 3.**
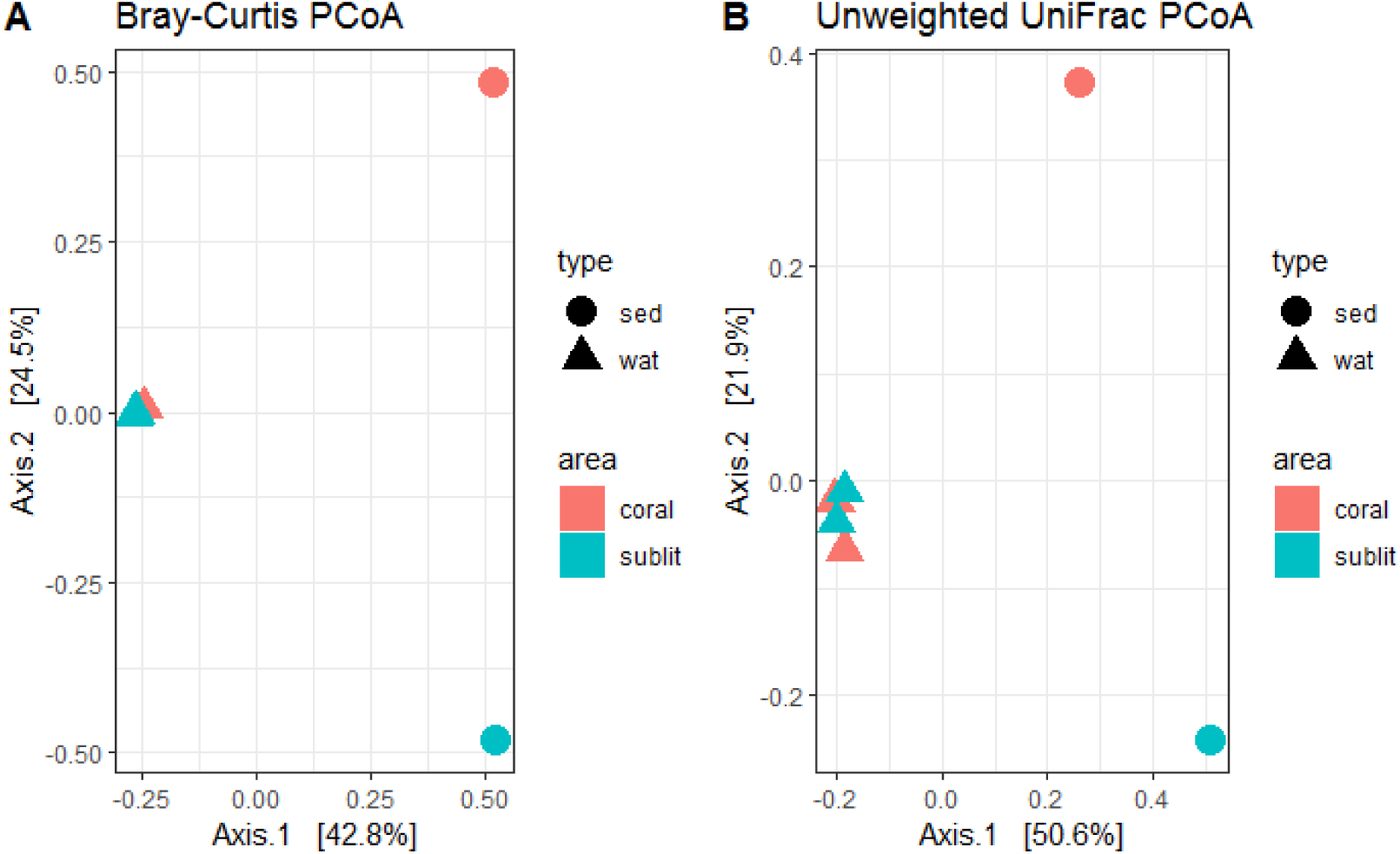
Principal coordinate analysis (PCoA) plots of (A) Bray–Curtis and (B) abundance-unweighted UniFrac distances. Shapes represent sample type: sed = sediments, wat = water, while colours represent sampling site: coral = coral reef, sublit = near-shore sublittoral.

**Fig. 4.**
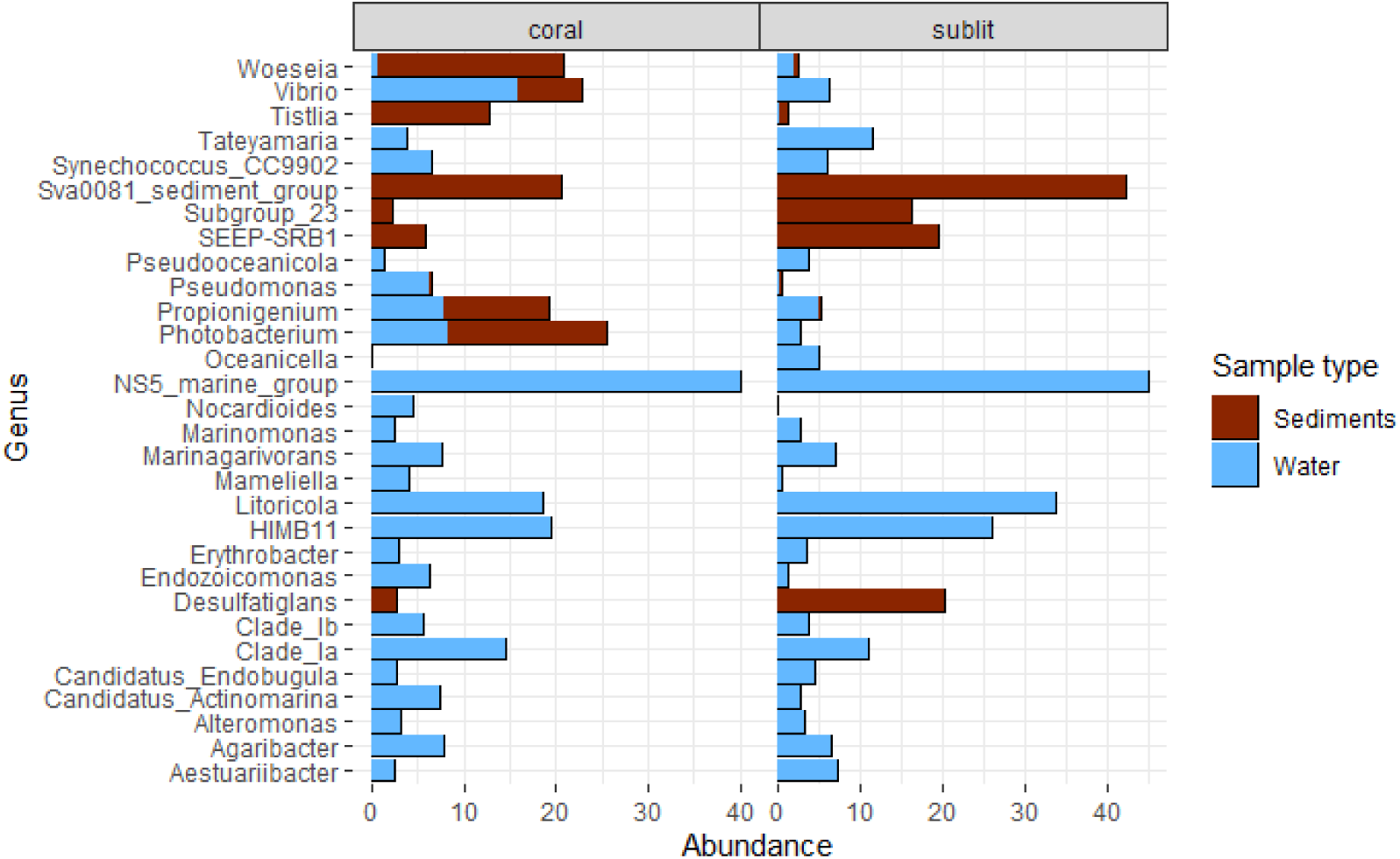
Relative proportion of the thirty most abundant bacterial genera. Each bar represents the genus distribution in sediment and water samples. Colour represents the sample type.

Over 96% of the RSVs were assigned to a known phylum with the proportion of assigned RSVs decreasing with lower taxonomic levels to ∼37% and <1% assignments at genus and species ranks respectively. A total of 41 microbial phyla were uncovered with majority of the RSVs being classified as Proteobacteria (∼64%), followed by Bacteroidetes (∼10%), Cyanobacteria (∼5%), Acidobacteria (∼4%), Planctomycetes (∼4%), Chloroflexi (∼3%), and Actinobacteria (∼2%). In water samples, ∼51% of the Proteobacteria phylum was made up of members of the *Gammaproteobacteria* class, while *Deltaproteobacteria* was dominant in sediment samples at ∼56%.

## Discussion

Characterizing marine microbial communities is critical not only for cataloguing a region’s genetic diversity but also for understanding the roles played by microorganisms in the fundamental processes of the marine ecosystems. The majority of studies to date estimating Indian Ocean biodiversity have focused on corals (Obura, 2012) and coral-associated eukaryotes (Kohn & Lloyd, 1973; Kohn & Nybakken, 1975; Van Der Elst et al., 2005). Of the three studies that assessed microbial diversity, one characterized only viruses (Williamson et al., 2012) while the two that evaluated bacterial communities (Sunagawa et al., 2015; Zinger et al., 2011) reported pooled findings for global oceans with little specific details, if at all, on the Indian Ocean. Moreover, only one of these studies sampled the coastal ocean and none within the coral reefs. To our knowledge this is the first description of water and sediment microbial population composition of WIO coral reef. Although limited sequencing replicates restricted the extent to which conclusive comparisons could be made, notable observations are cautiously discussed, and should be confirmed in the future through more robust studies.

Microbial communities in the sediments had higher diversity than in the water samples. This conclusion is reasonable as has been established through numerous studies of various aquatic ecosystems (Feng et al., 2009; Shulse et al, 2017; Sogin et al., 2006; Won et al, 2017). Higher microbial diversity in marine sediments is thought to be because sediments offer the surface for interaction of water column-derived organic matter with benthic microbial communities (Probandt et al., 2017). This is the case especially near the shore (Dyksma et al., 2016), which might explain why our near-shore sublittoral sediment samples had slightly higher numbers of observed ribotypes (889 RSVs) compared to samples from the coral reef (525 RSVs) located farther from the shore. The biogeographical separation between near-shore sublittoral and coral reef locations of our sampling sites could also explain why sediment microbial populations didn’t cluster as those from water samples. Clustering of coral reef and sublittoral water microbiota may be due to tidal mixing of water in these zones, a phenomenon that is less dynamic within sediments between the two zones.

The observed dominance of members of Proteobacteria, Bacteroidetes and Cyanobacteria phyla was expected considering findings from the global oceans (Sunagawa et al., 2015; Zinger et al., 2011) as well as previous studies of coral reefs and coastal associated niches (Curren & Leong, 2019; Feng et al., 2009; Godoy-Vitorino et al., 2017; Wegley et al., 2007). As expected, different classes of Proteobacteria phylum dominated water and sediment samples (Feng et al., 2009; Zinger et al., 2011): members of *Gammaproteobacteria* class were the most abundant in water samples whereas in the sediment samples *Deltaproteobacteria* were predominant.

Less than 1% of RSVs were assigned to a known species. Some of these species included *Marinobacter litoralis* which synthesizes extremozymes, such as halophilic lipase, with a wide range of biotechnological potential (Musa et al., 2019); *Pseudoalteromonas luteoviolacea* which produces a range of antibacterial, bacteriolytic, agarolytic and algicidal compounds as well as compounds that prevent settling of fouling organisms and promote survival of other marine organisms (Holmström & Kjelleberg, 1999); and *Pseudoalteromonas phenolica* which is known to produce specific antibacterial compounds against anti-methicillin-resistant *Staphylococcus aureus* (MRSA) (Isnansetyo & Kamei, 2003, 2009). We also identified *Ferrovibrio xuzhouensis*, a cyhalothrin-degrading bacterium (Song et al., 2015). These results confirm WIO’s biological richness, as well as the versatility and utility of environmental DNA (eDNA) methods in bioprospecting as has been rationalised before (Behzad et al., 2016; Synnes, 2007).

These results give the first glimpse of the WIO’s microbial diversity and showcase the potential of an eDNA metabarcoding approach as a survey and/or monitoring tool that may benefit marine resource conservation and management efforts.

## Acknowledgment

This work was made possible by logistical support of the Kuruwitu Conservation & Welfare Association and financial support from the Swedish Research Council. SW received a travel grant from the Western Indian Ocean Marine Science Association (WIOMSA) to facilitate data analysis.

